# A tissue specific atlas of gene promoter DNA methylation variability and the clinical value of its assessment

**DOI:** 10.1101/2022.02.04.479170

**Authors:** Ryan H Miller, Chad A Pollard, Kristin R Brogaard, Andrew C Olson, Larry I Lipshultz, Erica B Johnstone, Yetunde O Ibrahim, Jim M Hotaling, Enrique F Schisterman, Sunni L Mumford, Kenneth I Aston, Tim G Jenkins

## Abstract

**Background:** Complex diseases have multifactorial etiologies making clinically actionable diagnostic markers difficult to identify. Novel tools with higher diagnostic yield and utility in driving personalized care are needed.

**Methods:** We utilized Illumina methylation array data from over 2400 samples to assess DNA methylation patterns in 20 distinct cell types ranging from sperm to brain as well as various disease states. We generated a simple analysis pipeline for DNA methylation data that focuses on intra-individual methylation variability within gene promoters. The analysis is designed, not to identify single causative gene alterations but instead focuses on any movement away from “healthy” methylation. This approach identifies altered regulation across multiple genes in related pathways thus enabling us to detect shifts in gene regulatory activity associated with distinct tissues and phenotypes. We explored three distinct questions in our assessment. 1) Are patterns of epigenetic variability tissue specific? 2) Do diseased tissues exhibit altered variability patterns compared to normal tissue? 3) Can epigenetic variability be detected in complex disease?

**Results:** Unsupervised clustering analyses established that patterns of epigenetic variability are tissue specific and that these patterns are at least as predictive of tissue type as differential methylation analysis. We demonstrated the ability to use these patterns to differentiate between healthy and diseased tissue with unsupervised clustering even in cases of complex multifactorial diseases. We applied this method to the clinical use case of male infertility and found that men undergoing intrauterine insemination (IUI) with the lowest number of epigenetically dysregulated promoters in their sperm were almost twice as likely to father a child than men participating in IUI with the highest number of dysregulated promoters (p=0.011). We saw no significant difference in birth rates between groups of men with high and low numbers of dysregulated promoters undergoing in vitro fertilization (IVF), indicating IVF as a better treatment than IUI to achieve live birth in the presence of multi-pathway dysregulation in sperm.

**Conclusions:** This study demonstrates that patterns of epigenetic variability can differentiate between tissue types. While intuitive, this finding has never been demonstrated previously and suggests that specific epigenetic variability patterns may be used to predict phenotypic changes in disease states as these are, by definition, functional changes to cellular phenotypes. We demonstrate that the variability of gene regulatory marks are distinct between healthy and diseased tissue. This is particularly apparent at genes known to be important to cell function of the tissue of interest. While in some cases these regional alterations can be seen across the entire genome, more often the regulatory alterations that define a pathological phenotype are restricted to genes of known importance to a particular tissue. Importantly, in the case of sperm, we found that these patterns of variability did have utility in predicting infertile patients who would conceive through intrauterine insemination (IUI). We would propose that this discriminatory ability is due to the fact that the signature can be assessed in an n-of-1 context and that the patterns of variability identify any shift away from regulatory normalcy in pathways known to be impactful in the tissue of interest, and not only assessing the presence or absence of rare genetic variants. While the data presented here are encouraging, more work needs to be performed in other tissues to determine when, and in what context, these findings could be clinically actionable.

## Introduction

In 2003, one of the most profound efforts ever undertaken in the biological sciences, the Human Genome Project, was completed[1]. At the time there was a great deal of hope that unlocking the genetic code was the key to diagnosing and treating the vast majority of diseases. While the discoveries made have been of great interest to many and have opened the door for important genetic and epigenetic findings, clinically meaningful impacts remain elusive for many diseases. In large part, this is due to the complex, multifactorial nature of most disease processes, with etiologies resulting from a constellation of genetic, epigenetic, and environmental perturbations. As a result, novel approaches to our analysis of genetic and epigenetic data are needed to identify clinically actionable predictors of disease or disease progression.

In hindsight, the lack of clinical efficacy for many epigenetic and genetic findings is not surprising. While it is common to identify a genetic or epigenetic association to a given disease, those associations occur so rarely in a population, or are only one of many alterations required to generate a pathological phenotype, that it is unreasonable to produce a highly predictive screening test capable of impacting clinical care. Importantly, while the causative factors of complex disease are multifactorial and highly variable, the tissue and cellular phenotypes that occur as a result of the disease are likely to be more conserved. Therefore, novel diagnostic approaches should focus, not on single genes or independent modifications associated with a pathology, but on a holistic screen of alterations to gene regulatory activity at genes specifically important to the affected tissue.

Certain data types are ideal for analyses focused on perturbations to gene regulatory networks. DNA methylation is of particular interest in this effort. Because a tissue’s phenotype is defined by gene activity, and gene activity is controlled (at least in part) by epigenetic marks, it is intuitive that marks such as DNA methylation are effectively the fingerprint of cell and tissue type. A logical extension of this idea is the fact that DNA methylation and other epigenetic marks are altered in diseases such as cancer[2,3] and type 2 diabetes[4,5] not because they are causative features of the disease, but because the diseases induce epigenetic modifications to achieve a perturbed phenotype. Further, unlike the fairly static nature of the genome, DNA methylation is a dynamic biomarker affected by a host of factors such as age[6,7] and various modifiers including obesity[8,9], exercise[10,11], and environment[12]. Despite the evidence that clearly suggests that DNA methylation is tied to complex disease states, we need to use novel analytical approaches to realize its full potential.

Herein, we present a novel approach to DNA methylation analysis and an assessment of its efficacy as a clinical predictor. One recent study used a similar method specifically for women’s cancers and found significant utility of the approach[13]. In our case, we describe a simple analysis of promoter DNA methylation variability that allows us to assess gene regulatory networks in a novel way. We provide evidence using DNA methylation array data from over 2400 samples and 20 cell types that our analysis can be used to identify and assess the genes most tightly regulated in specific cell types and that these patterns are highly tissue-specific. We then show the utility of this approach in an n-of-1 analysis to demonstrate that there are gene regulatory network perturbations common among individuals who suffer from specific pathologies in many different tissues. Lastly, we demonstrate in sperm the potential clinical value of this approach.

## Methods

### Data Collection

Several publicly available datasets were used in this study. Infinium HumanMethylation450 Bead Chip data was obtained for tumor and healthy tissue samples (n=494) from The Cancer Genome Atlas (TCGA) Program as compiled by the University of California Santa Cruz Xena Functional Genomics Explorer[14] (https://xenabrowser.net/datapages/). Infinium HumanMethylation450 Bead Chip data for CD4+ T cell (n=11), CD8+ T cell (n=14), Alzheimer’s disease and control brain (n=190), lung (n=6), liver (n=26), and skin methylation data (n=18) from healthy and diseased individuals were accessed from the NIH Gene Expression Omnibus (GSE130029, GSE130030, GSE66351, GSE51077, GSE61258, GSE115797, respectively).

Sperm Infinium HumanMethylation450 Bead Chip data from fertile sperm donors as well as patients undergoing in vitro fertilization (IVF) (n=166) was used from a previously published single-site study by Aston, et al[15] as well the sperm Infinium MethylationEPIC Array data from a clinical multi-site study of patients being seen by physicians for fertility care (n=1311) as published by Jenkins et al[16].

### Sample Collection

Semen samples were procured from University of Utah Andrology department from consented patients undergoing fertility care (n=60), as well as two independent fertile sperm donor cohorts (n=59). Semen samples from consented patients seeking fertility care were also procured from the Urology Department at Baylor College of Medicine (n=49).

### Sample Preparation

For all semen samples, somatic cell lysis, sperm isolation, DNA extraction, and bisulfite conversion were performed as described by Aston, et al[15]. The bisulfite converted sperm DNA was hybridized to Illumina Infinium HumanMethylationEPIC microarrays and ran as recommended by the manufacturer (Illumina) at Infinity BiologiX.

### Data Preprocessing

Figure 1 contains a flow chart of data processing and statistical analysis. The raw methylation array data from the sperm, neuron, glia, skin, CD4+ T cell, and CD8+ T cell samples were preprocessed using the minfi R package[17] using SWAN normalization to produce beta and m-values for each cytosine-guanine dinucleotide (CpG). Density plots of the beta values of each sample were examined to ensure the distribution of beta values followed a bimodal distribution with prominent peaks between 0.0-0.2 and 0.8-1.0 and flat valleys from 0.2-0.8. Any samples not following this distribution were removed and the remaining samples were renormalized. Beta values are described as (methylated probe intensity / [methylated + unmethylated probe intensity + 100]) and range from 0-1 with values around 0 being unmethylated and values around 1 being methylated. M-values are described as (log(methylated probe intensity / unmethylated probe intensity) and are useful measures of methylation to prevent bias arising from heteroscedasticity seen when analyzing beta values[18] (Supplementary Figure S1, Additional File 1).

**Figure 1:**
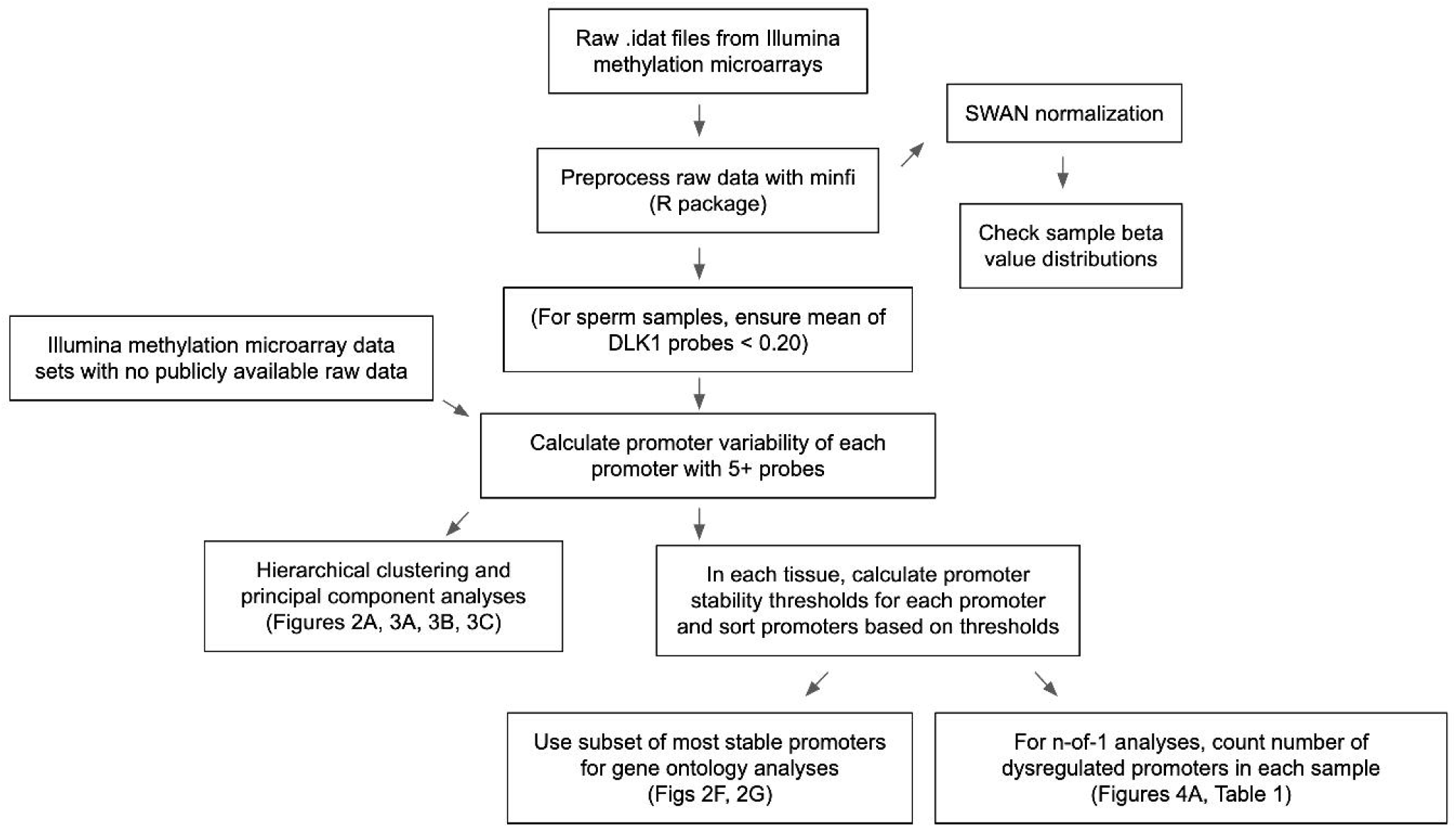
Data processing and statistical analysis workflow. Processing and analysis of Illumina’s Infinium HumanMethylation450 and HumanMethylationEPIC array data from multiple tissue types to derive promoter variability and promoter stability thresholds and analyze their relationships among tissue types and between healthy and diseased tissues.

For data processing of sperm samples, we removed any sperm samples from analysis that did not have a mean methylation value less than 0.20 of all the CpG beta values in the differentially methylated region of DLK1 as described by Jenkins, et al[19] (chr14:101,191,893-101,192,913, GRCh37). According to Jenkins et al, the methylation states of the probes in this region are a good discriminator between sperm and somatic cells and this procedure ensured analyses were only performed on samples containing sperm DNA methylation and not contaminating somatic cell DNA methylation.

Raw data for the TCGA datasets as assembled on the UCSC Xena platform and the lung and liver datasets (GSE51077 and GSE61258, respectively) were not available, so the available beta values were used. These beta values were logit-transformed to obtain the m-values for these samples.

### Statistical analysis

We define a given gene promoter as the genomic region one kilobase upstream and one kilobase downstream from the transcription start site of a given gene. A gene promoter needed to contain five or more methylation array probes to be used in any downstream analysis. Gene methylation promoter variability (or “promoter variability”) is defined as the standard deviation of the m-values of the methylation array probes present in a defined promoter region (Supplementary Figure S2, Additional File 1).

Hierarchical clustering was performed on all promoter variability values of samples from various tissue types using the R software package ‘pheatmap’ (R version 4.0.3) with default parameters. In cases where more than 20 samples existed for a given tissue, 20 samples were randomly selected for inclusion in the clustering analysis to give a more uniform number of samples per tissue type. Principal component analyses were performed on all promoter variability values using the ‘sklearn’ library in Python (Python version 3.7.3).

We found the most epigenetically stable promoters of a given tissue type by identifying the promoters with the lowest levels of variability in healthy samples of that tissue type. We did this by first calculating a stability threshold for each promoter in a given tissue. A promoter stability threshold represents the highest level of variability we expect to see in a given promoter of a healthy sample of a given tissue (Supplementary Figure S2, Additional File 1). Then, the promoters were rank ordered by the stability threshold values in ascending order. For the analyses comparing promoters across tissue types (Figures 2B, 2C, 2F, 2G), we defined the most stable promoters as the top first percentile of promoters with the lowest stability thresholds in healthy samples of the given tissue. The most stable promoters for the sperm n-of-1 analyses were defined as the top 10th percentile of promoters with the lowest stability thresholds in fertile sperm donors.

**Figure 2:**
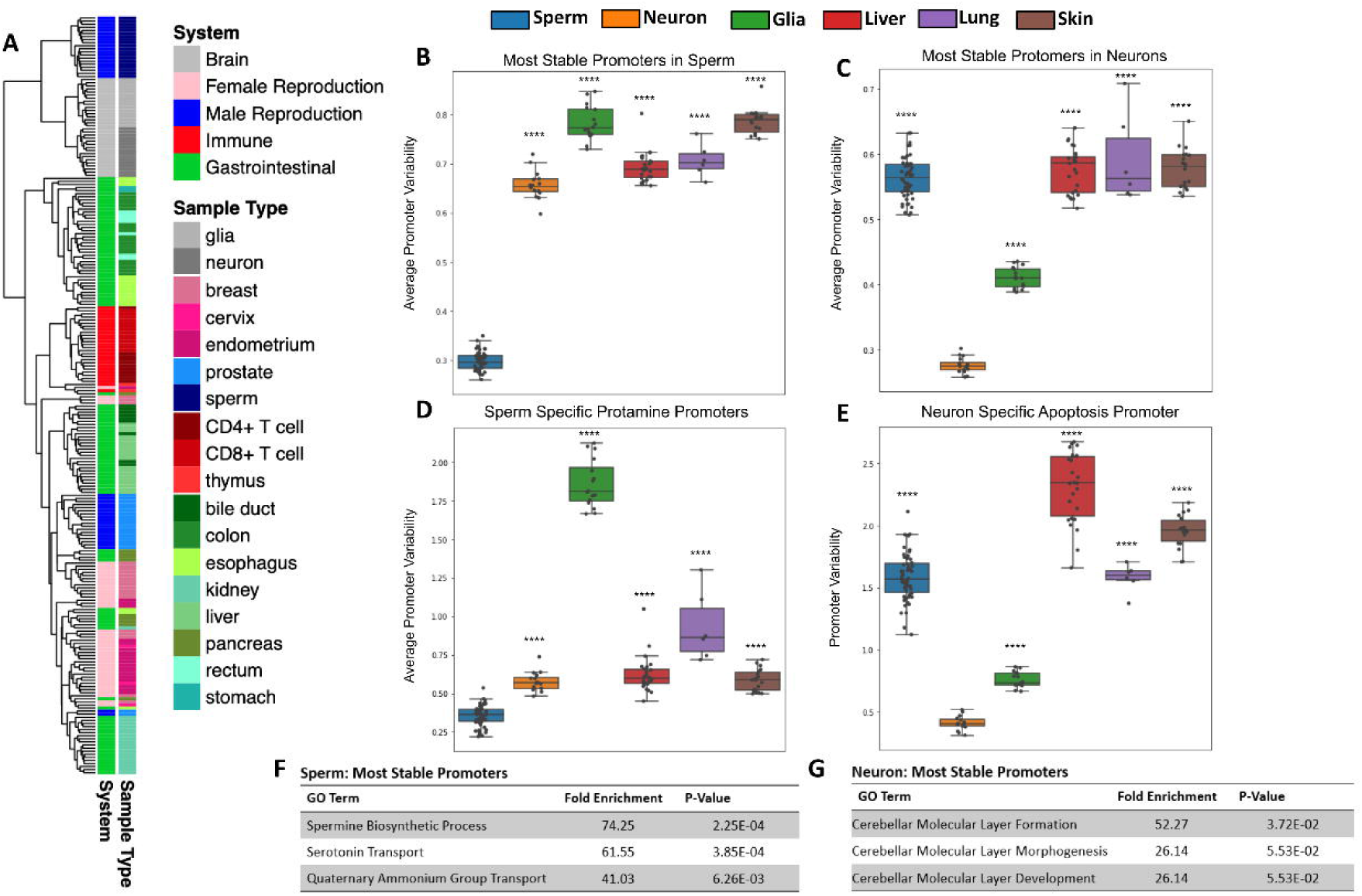
Tissues have unique patterns of gene methylation promoter variability. **A)** Hierarchical clustering of promoter variability of 18 sample types representing five biological systems. **B)** and **C)** Average promoter variability of 6 distinct cell types in the most stable promoters (top 1st percentile) in sperm and neurons, respectively. One dot represents one sample, and boxplots are overlaid to show the distribution of average promoter variability of each tissue. All p-values comparing methylation variance between sperm and neuron to other tissues types were ≤ 5.16E-14. **D)** Average protomer variability in the 6 cell types of three sperm-specific protamine promoters. **E)** Promoter variability from one neuron specific apoptosis promoter in the 6 cell types. All p-values for panels D and E when comparing sperm and neurons to other cell types were ≤ 9.99E-17. **F)** and **G)** Gene ontology enrichment of the most stable promoters for sperm and neurons, respectively.

Sperm n-of-1 analyses were performed by finding the most stable promoters in a cohort of fertile sperm donors and counting the number of dysregulated promoters in each sample. We defined a dysregulated promoter as a promoter that fell above the corresponding variability threshold. Samples with the lowest number of dysregulated promoters are most similar to healthy controls.

Statistical differences in the pregnancy and live birth rates of men undergoing intrauterine insemination (IUI) and in vitro fertilization (IVF) with the least and most dysregulated promoters were calculated with two-sided t-tests. The men undergoing IUI had been through an average of 2.5 IUI attempts.

Tissue-specific gene ontology enrichment analyses were performed by running the PANTHER Overrepresentation Test (http://pantherdb.org/webservices/go/overrep.jsp) on the gene names of the first percentile of most stable promoters in a given tissue. Each test was run using a background gene set that consisted of all genes with promoters containing five or more methylation array probes.

## Results

### Tissues have unique methylation variability signatures

Using microarray DNA methylation data, we explored the differences in gene promoter methylation variability of various healthy tissues. Unsupervised clustering of all gene promoter variability values showed tissue specificity and also revealed similarities among related tissues, Figure 2A. For example, gastrointestinal tissue samples such as those from the esophagus, stomach, colon and rectum, clustered closely together. We likewise saw clustering of samples associated with the immune system (CD4+ T cells, CD8+ T cells, thymus), female reproduction (endometrium, cervix), and brain (glia, neurons).

### Tissues are regulated at tissue-specific biological pathways

We sought to identify how promoter variability differs among various tissue types. We identified the most stable promoters in sperm and assessed the average methylation variability values for these promoters in many samples across several tissue types as seen in Figure 2B. At promoters indicated as most stable in the male germline, sperm samples have significantly lower mean values than other tissue samples. Gene ontology analysis of these sperm promoters show significant enrichment for sperm-related biological processes (Figure 2F). We also looked at the mean of the promoter variability values of the known sperm-related genes protamine 1 (PRM1), protamine 2 (PRM2), and protamine 3 (PRM3) which are genes expressed exclusively in sperm and replace the majority of histones to achieve extreme nuclear compaction in this specialized cell[20]. As expected, sperm samples displayed significantly less variability in these promoters than other tissues (Figure 2D). These same analyses were performed for the most stable promoters in neurons (Figures 2C, 2E) and a known neuron-specific gene, CASP8 (Figure 2G) with similar results. Additional File 1 (Supplementary Figure S4, Additional File 1) contains the results of these same analyses performed for several other tissue types. It is important to note that while the most stable promoters in a given tissue are generally characterized by very low promoter variability in samples from the given tissue, these promoters all have varying degrees of absolute methylation (hypo, mid, or hyper-methylation), a feature missed by traditional differentially methylated region (DMR) analyses (Supplementary Figure S5, Additional File 1). It is also conceivable that this method can help overcome technical biases inherent to methylation microarrays such as batch effects.

### Methylation variability can differentiate between healthy and diseased tissue

In addition to distinguishing between tissue types, analysis of promoter methylation variability can enable the differentiation of diseased and healthy tissue samples of the same tissue type. One notable example is the ability to distinguish between tumor and healthy tissue based on promoter variability signatures. Figure 3A depicts the first two principal components of promoter variability values for colon primary tumor tissue and healthy colon tissue. The healthy colon tissue samples appear to be tightly clustered together, whereas the tumor samples are widely distributed throughout the plot. Figure 3B shows the difference in promoter variability between psoriatic skin lesions and adjacent healthy skin samples from the same individuals. Figure 3C shows a principal component analysis of neurons, glial cells, as well as bulk cell samples from postmortem brain tissue of individuals with Alzheimer’s disease and controls. The plot shows clear separation among the neurons, glial cells, and bulk cell samples indicating a difference in promoter variability among different cell types in the same tissue. There is also separation between control and Alzheimer’s disease samples in neuron and glial cell samples but such separation is not apparent among the bulk cell samples which may suggest subtle differences in promoter variability might be more apparent when samples are sorted for individual cell types.

**Figure 3:**
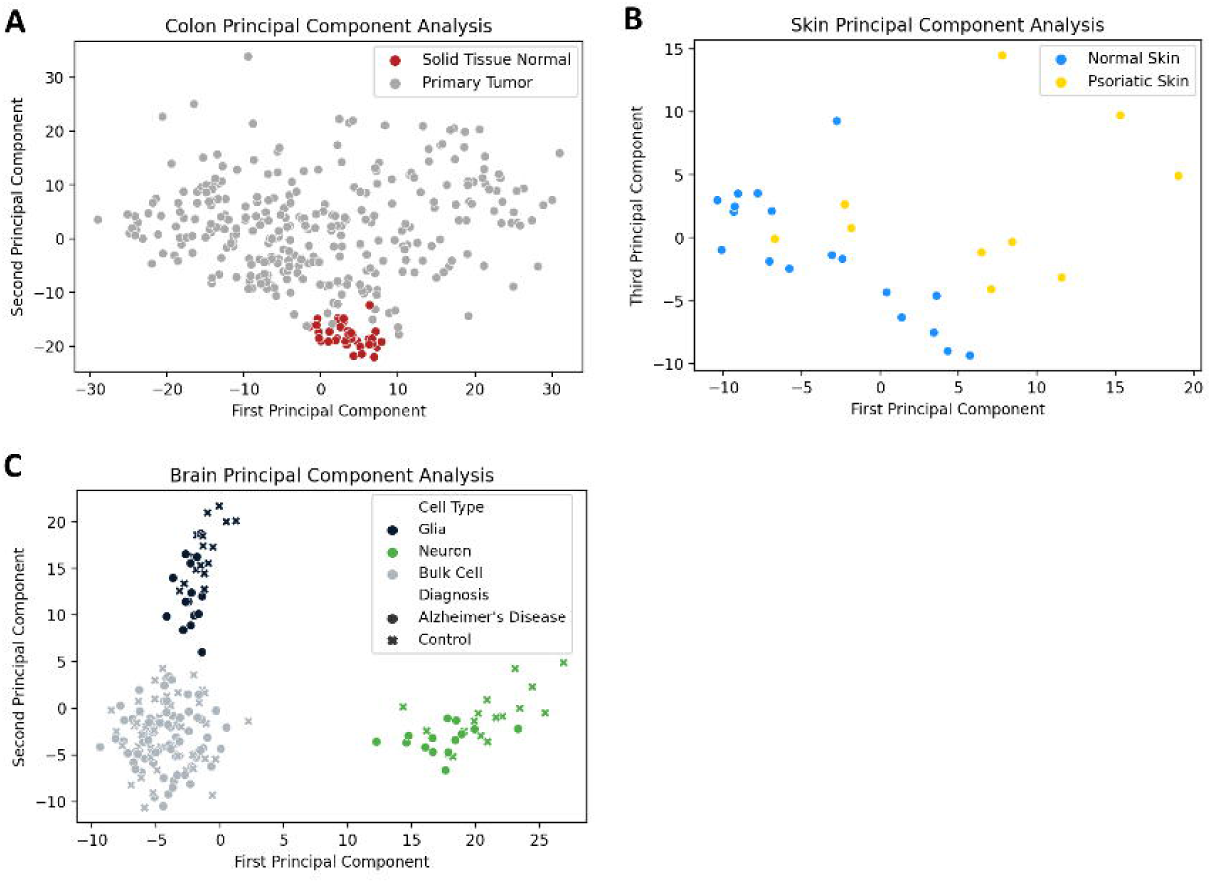
Diseased tissue samples have unique patterns of gene methylation promoter variability compared to healthy. **A)** Principal component analysis of promoter variability values from primary colon tumors and normal colon tissue. **B)** Principal component analysis of promoter variability values from matched psoriatic lesion and healthy skin samples. **C)** Principal component analysis of promoter variability among neurons, glial cells, and bulk cells from postmortem brains of individuals with Alzheimer’s disease as well as controls.

### Methylation variability of sperm can identify a subset of men with male factor infertility

We performed n-of-1 analyses on over 1500 sperm samples from fertile sperm donors as well as men being treated for male factor infertility. Figure 4A shows there was a significantly higher number of dysregulated promoters in men being treated for male factor infertility compared to fertile sperm donors. Supplementary Figure S8 (Additional File 1) shows the number of dysregulated promoters in multiple cohorts of sperm samples, including the sperm donor cohort used to find the most stable promoters in sperm as well as the stability thresholds. To give a visual explanation of promoter methylation, Figure 4B depicts the promoter variability values (red dots) of the most stable sperm promoters and the corresponding stability thresholds for these promoters (black line) in an individual fertile sperm donor sample as well as a patient being treated for male factor infertility (Figure 4C). It is clear that the patient being treated for male factor infertility has many more dysregulated promoters than the fertile sperm donor. This suggests that male factor infertility may be more related to a global shift in methylation variability at promoters important for sperm cells rather than single nucleotide changes or epimutations.

**Figure 4.**
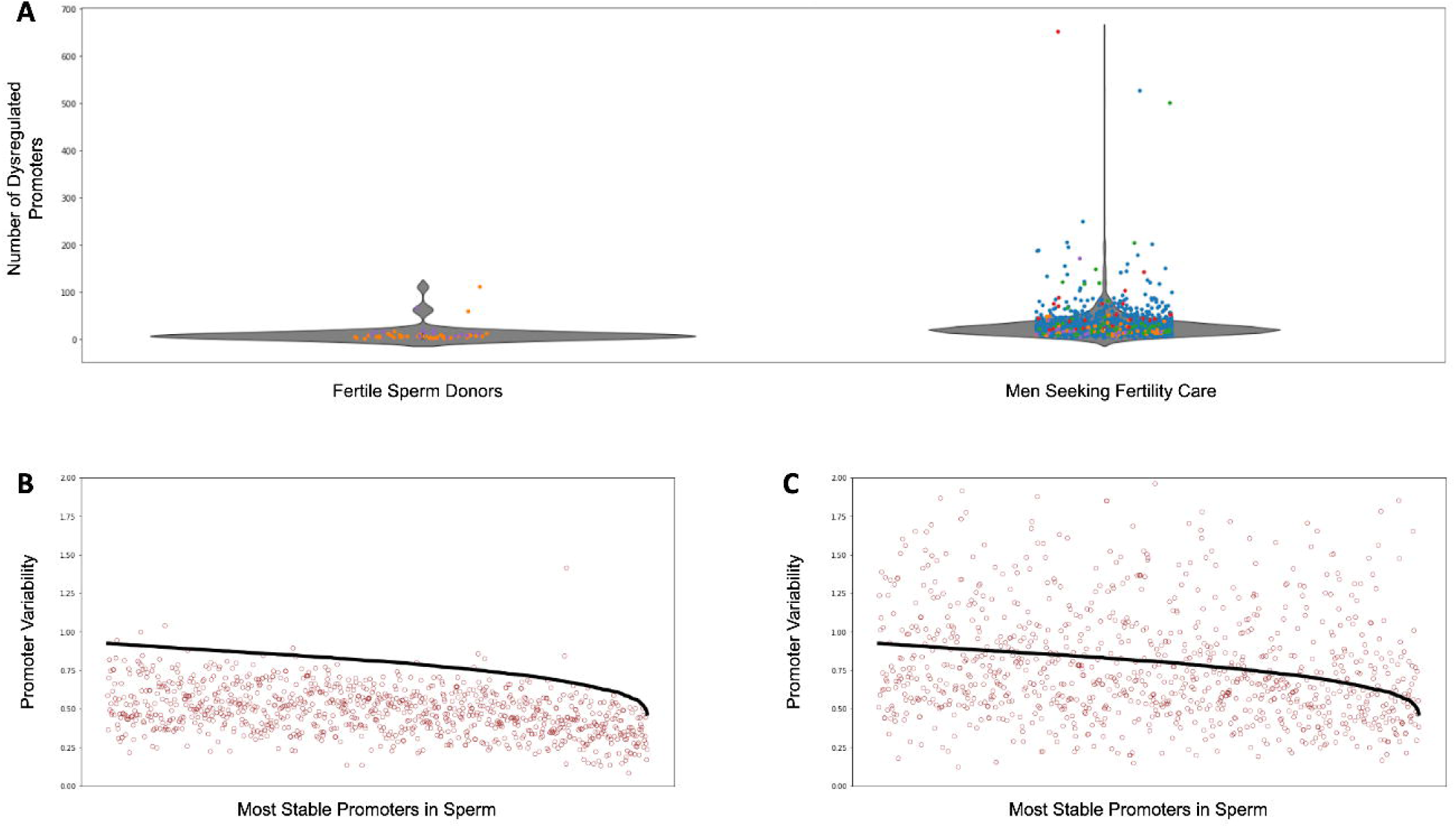
Dysregulated promoters are enriched in men seeking fertility care compared to fertile controls. **A)** Dysregulated promoters in samples from five independent studies. The most stable sperm promoters and corresponding stability thresholds were calculated from a cohort of fertile sperm donor samples. Circle colors represent the following: blue - Jenkins et al[16], green - Baylor College of Medicine, purple - fertile sperm donors, Red and Orange - University of Utah Andrology two independent collections. **B)** Promoter variability at the most stable sperm promoters in a single fertile donor sperm sample (red dots). The stability threshold for these promoters are shown in black. A red dot above the black line indicates a dysregulated promoter. **C)** Same analysis as in B) but for a single patient being treated for male factor infertility.

We then looked at the relationship between dysregulated promoters and pregnancy and live pregnancy rates in 1311 individuals being seen by a physician for infertility care. Table 1A shows that in men undergoing intrauterine insemination (IUI), those with least number of dysregulated promoters (lowest 10th percentile of patients) had significantly higher pregnancy and live birth rates than IUI patients with the most number of dysregulated promoters (top 10th percentile of patients). However, we saw no difference in pregnancy and live birth rates when comparing men undergoing in vitro fertilization (IVF) with the least and most number of dysregulated promoters, suggesting IVF should be the preferred fertility treatment option for men with high levels of methylation dysregulation.

**Table 1.** Pregnancy and live birth rates in patients with the least and most dysregulated promoters. **A)** Pregnancy and live birth rates from male patients undergoing on average 2.5 intrauterine insemination (IUI) cycles (N=528). **B)** Pregnancy and live birth rates from male patients undergoing IVF (N=234). For each patient’s sperm sample, the number of dysregulated promoters were counted. Pregnancy and live birth rates were compared between the patient cohort with the top 10th percentile bottom 10th percentile of dysregulated promoters.

| Patient Cohort | Pregnancy Rate from IUI | Live Birth Rate from IUI |
| --- | --- | --- |
| Patients with the least number of dysregulated promoters (bottom 10th percentile, n=54) | 48.1% | 40.7% |
| Patients with the most number of dysregulated promoters (top 10th percentile n=50) | 26.0% | 18.0% |
|  | p=0.019 | p=0.011 |

| Patient Cohort | Pregnancy Rate from IVF | Live Birth Rate from IVF |
| --- | --- | --- |
| Patients with the least number of dysregulated promoters (bottom 10th percentile, n=36) | 75.0% | 61.1% |
| Patients with the most number of dysregulated promoters (top 10th percentile, n=23) | 73.9% | 60.9% |
|  | p=0.927 | p=0.986 |

## Discussion

Here, we introduce a novel method which assesses gene promoter DNA methylation variability to identify highly regulated genes in multiple tissue types and how these can be impacted in various disease states. Hierarchical clustering of gene promoter variability from many tissues demonstrates how unique these patterns are in different tissues and how these patterns remain largely consistent in related, but distinct tissues. For example, numerous tissues from the gastrointestinal tract cluster together as do tissues important to the function of the immune system.

We also found that the most stable promoters in a given tissue have significantly lower methylation variability than the same promoters in other tissues highlighting the importance of unique genes and gene networks to any given tissue’s function. Importantly, when assessing DNA methylation variability within the same tissue type, we were also able to visualize differences between healthy and diseased tissues such as in cancer, psoriasis, and Alzheimer’s disease.

To highlight the potential clinical impact of the assessment of promoter level DNA methylation variability, we examined the pattern’s utility in an assessment of male factor infertility and found that men being seen by a physician for infertility had much higher levels of dysregulated promoters. In addition, men undergoing IUI treatments with the most number of dysregulated promoters had significantly lower pregnancy and live birth rates compared to men undergoing IUI treatments with the least number of dysregulated promoters. However, this stark difference in pregnancy and live birth rates was not seen between men with the highest and lowest levels of dysregulated promoters in men undergoing IVF. This finding has great clinical significance because it suggests that if a man is struggling with infertility and has a high level of promoter dysregulation, he has much better odds at having a child after undergoing IVF than if he simply went through multiple rounds of IUI.

Taken together, these data suggest that promoter DNA methylation variability is an excellent indicator of tissue type. This is likely due to the fact that the analysis of variability is able to successfully detect genes that are the most tightly regulated (via DNA methylation in this case) in any given tissue. The assumption is that these genes must play an important role in cell function unique to each tissue. This is supported by our data that demonstrated that promoter DNA methylation variability is increased on average in abnormal tissues when compared to normal tissue. This was particularly apparent in our assessment of sperm DNA methylation variability patterns. While much work still needs to be performed, these patterns are promising and may prove to have significant utility in clinical decision making.

Of additional value in our assessment of this approach and its translational capacity is the ability to perform these analyses in an n-of-1 context. Specifically, because we are assessing intra-promoter variability within a single individual, we are able to reliably assess variability in a single individual with limited concerns of batch effects which would require normalization. While this has yet to be fully vetted in the current work, it does appear that these analyses, if proven to have high predictive power, could be translated to a diagnostic tool.

While we did perform one of the largest analyses to date in terms of tissue types and sample numbers, many questions still remain to be answered. Among the most critical is the utility of this analysis in many different tissues and disease states. Because we were only able to perform a deep analysis in sperm due to the large numbers of samples, similar work needs to be done in other tissues to determine if this analysis can provide clinically actionable information. Even in our sperm study, this work needs to be replicated in independent cohorts to determine its efficacy. One area of concern for this type of analysis is the fact that not all tissues are able to be as easily purified as sperm. In fact, we found that when looking at bulk tissues in the brain, we lost any ability to discriminate between healthy and diseased tissue. Thus, future work needs to focus on as pure of tissue as possible to yield meaningful results. This could create technical challenges that make the utilization of this analysis untenable in some applications. Despite this, there will certainly be tissues and diseases in which the assessment of promoter DNA methylation variability will be feasible.

These findings provide a novel means to define which genes each cell and tissue type tightly regulate to ensure their unique phenotype and function. Because these signals have potential utility in both the a basic understanding of tissue specific epigenetic patterns and in the clinical assessment of diseased tissues, as well as the prediction of outcomes, this work provides important foundational findings upon which tissue and disease specific assessments can be constructed in the future. While much work remains to determine clinical actionability for various applications, the results here are encouraging and may offer another tool with which we can assess the health of tissues and, importantly, predict the outcomes from various clinical interventions.

## Availability of data and materials

The data from this study are available from the corresponding author upon reasonable request.

## Supplementary Information Figure Legends

### Additional File 1: Supplementary figures and tables

**Figure S1**. Heteroscedasticity of beta values in a sperm donor sample. **Figure S2**. Variability equations. **Figure S3**. Hierarchical clustering of different control tissues using beta values **Figure S4**. Average promoter variability at tissue-specific promoters. **Figure S5**. Example of promoters with low methylation variability but varying levels of methylation. **Figure S6**. Principal component analysis of diseased and control samples. **Figure S7**. Hierarchical clustering of diseased and control tissue samples. **Figure S8**. N-of-1 dysregulated promoter analysis of sperm samples from multiple cohorts. **Table S1**. Pregnancy/Birth rates in men with normal sperm concentration (≥ 15 million sperm / mL). **Table S2**. Statistics from men with the least and most number of dysregulated promoters. **Table S3**. Ranking by sperm concentration.

## Acknowledgements

The results shown here are in part based on data generated by the TCGA Research Network: https://www.cancer.gov/tcga.

## Author information

Ryan H Miller and Chad Pollard contributed equally to this work. Tim G Jenkins and Kenneth I Aston contributed equally to this work.

## Contributions

TGJ and KIA conceived and designed the study. RHM, CAP, and KB performed all analyses. RHM, CAP, KRB, KIA, and TGJ wrote the manuscript. All authors read and approved the final manuscript.

## Ethics Declarations

### Ethics approval and consent to participate

All research and sample procurement was conducted in accordance with the regulations and guidelines established by the Institutional Review Boards of University of Utah (IRB_00012049, University of Utah) and Baylor College of Medicine (H-50666, Baylor College of Medicine).

Consent was obtained from all participants prior to their donation of tissue samples. All other data sets were publicly available.

### Consent for publication

Not Applicable

## Supplementary Information

Additional File 1

Supplementary figures and tables

