## Additional File 1 for "A tissue specific atlas of gene promoter DNA methylation variability and the clinical value of its assessment"


Figure S1 | Heteroscedasticity of beta values in a sperm donor sample 2

Figure S2 | Variability equations 3

Figure S3 | Hierarchical clustering of different control tissues using beta values 4

Figure S4 | Average promoter variability at tissue-specific promoters 5

Figure S5 | Example of promoters with low methylation variability but varying levels of methylation 6

Figure S6 | Principal component analysis of diseased and control samples 7

Figure S7 | Hierarchical clustering of diseased and control tissue samples 8

Figure S8 | N-of-1 dysregulated promoter analysis of sperm samples from multiple cohorts 9

Table S1 | Pregnancy/Birth rates in men with normal sperm concentration (≥ 15 million sperm / mL) 10

Table S2 | Statistics from men with low and high number of dysregulated promoters 10

Table S3 | Ranking by sperm concentration 10

**Figure S1 | Heteroscedasticity of beta values in a sperm donor sample**
(A) Distribution of mean promoter methylation of the 100 most stable promoters in sperm which were found by calculating the variability value of the beta values of all probes in a promoter region. (B) Distribution of mean promoter methylation of the 100 most stable promoters in sperm which were found by calculating the variability value of the m-values of all probes in a promoter region.

A

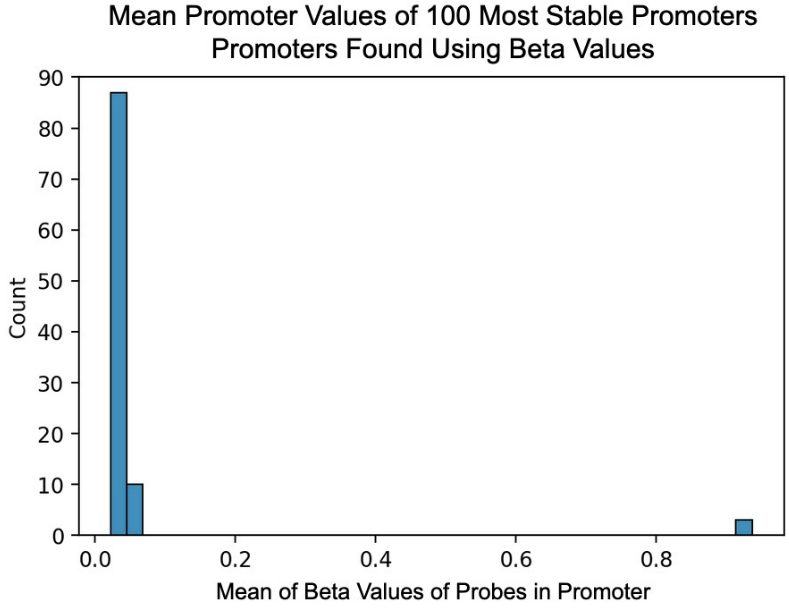


B

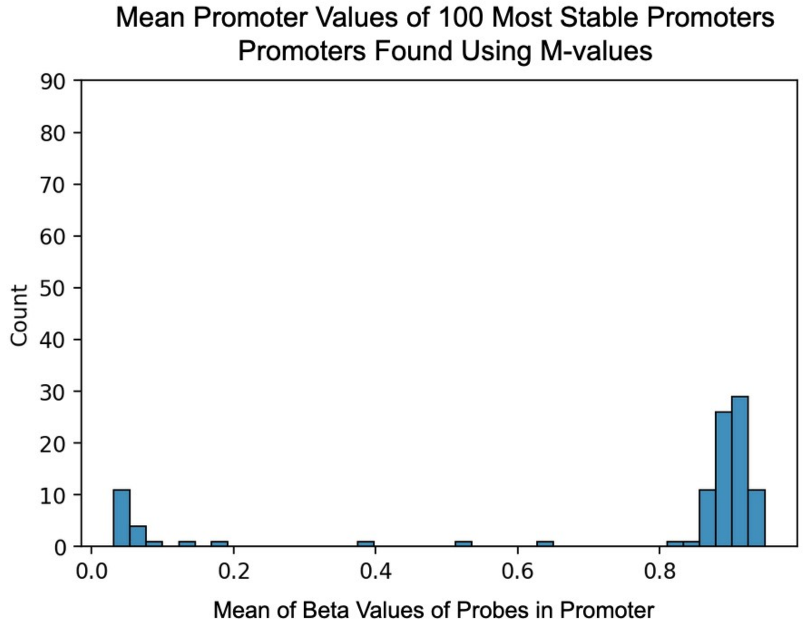

**Figure S2 | Variability equations**

(A) Equation for calculating the variability value (or standard deviation) of a given promoter in a sample; $\sigma$ = gene promoter variability value, $x_1$ = m-value of a given methylation array probe in a given promoter, $\mu$ = mean of probe m-values in given promoter (B) Equation to calculate the promoter variability threshold for a given tissue. $\Theta$ = promoter variability threshold for a given tissue, $\sigma_1$ = promoter variability value of a sample in a given cohort at a given promoter, $\mu$ = mean of probe m-values in a given promoter

A

$$\sigma = \sqrt{\frac{\sum |x_1 - \mu|^2}{N}}$$

B

$$\theta = \frac{\sum \sigma_1}{N} + 3\sqrt{\frac{\sum |\sigma_1 - \mu|^2}{N}}$$

**Figure S3 | Hierarchical clustering of different control tissues using beta values**
This plot was created by performing hierarchical clustering on all promoter mean values of the beta values of probes in a given promoter.

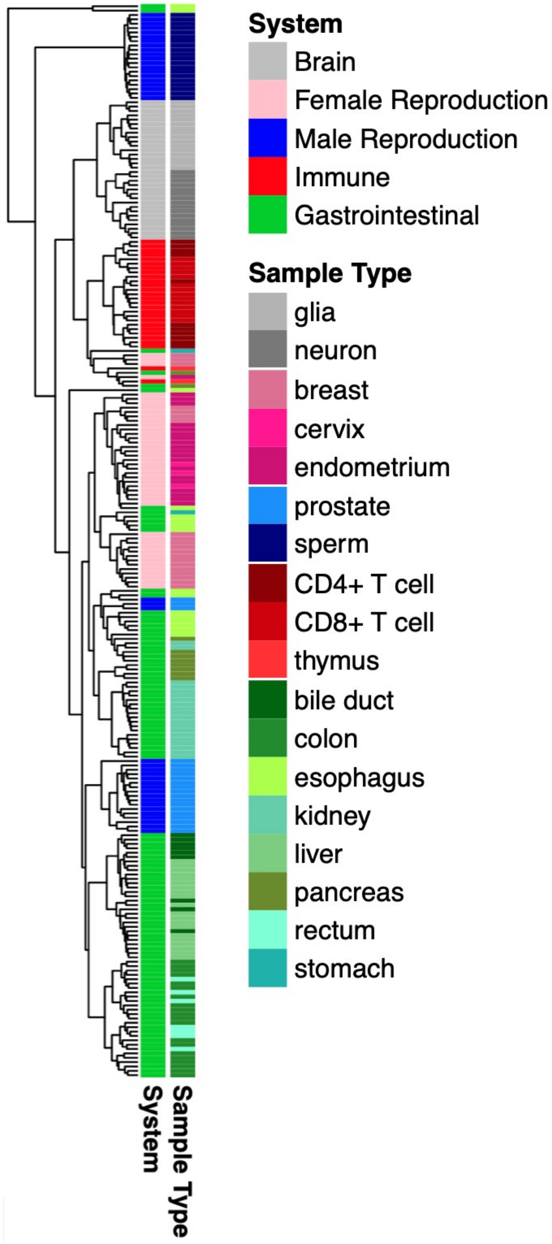

**Figure S4 | Average promoter variability at tissue-specific promoters**
(A) Average promoter variability values of numerous samples across several tissues at most stable promoters in control lung tissue. (B) Average promoter variability values of numerous samples across several tissues at most stable promoters in control skin tissue. (C) Average promoter variability values of numerous samples across several tissues at most stable promoters in control liver tissue.

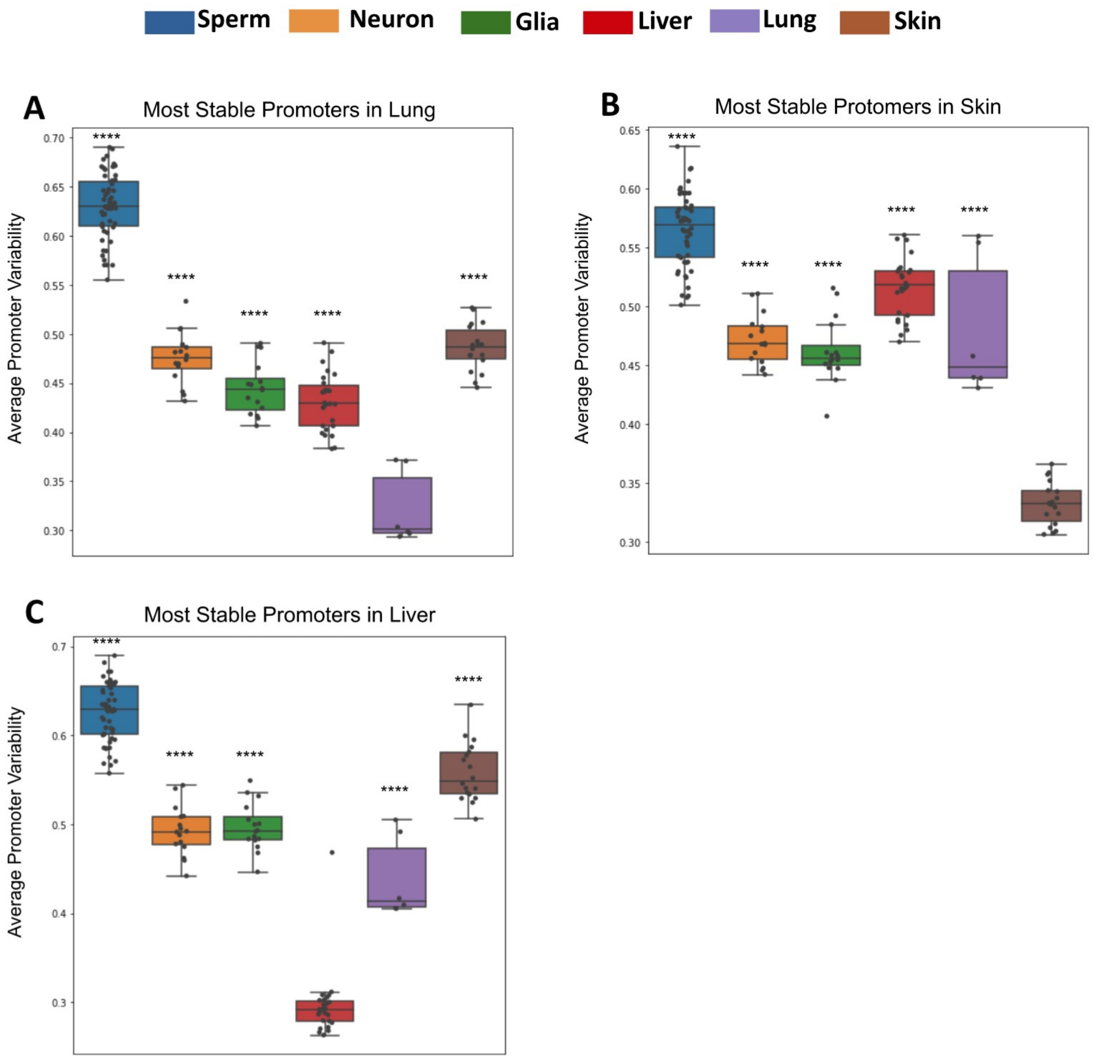

**Figure S5 | Example of promoters with low methylation variability but varying levels of methylation**

(A) Boxplot of methylation values of 31 sperm donor samples at 3 gene promoters with low methylation variability (B) Dotplot of methylation values of 1 sperm donor sample at 3 gene promoters with low methylation variability

A

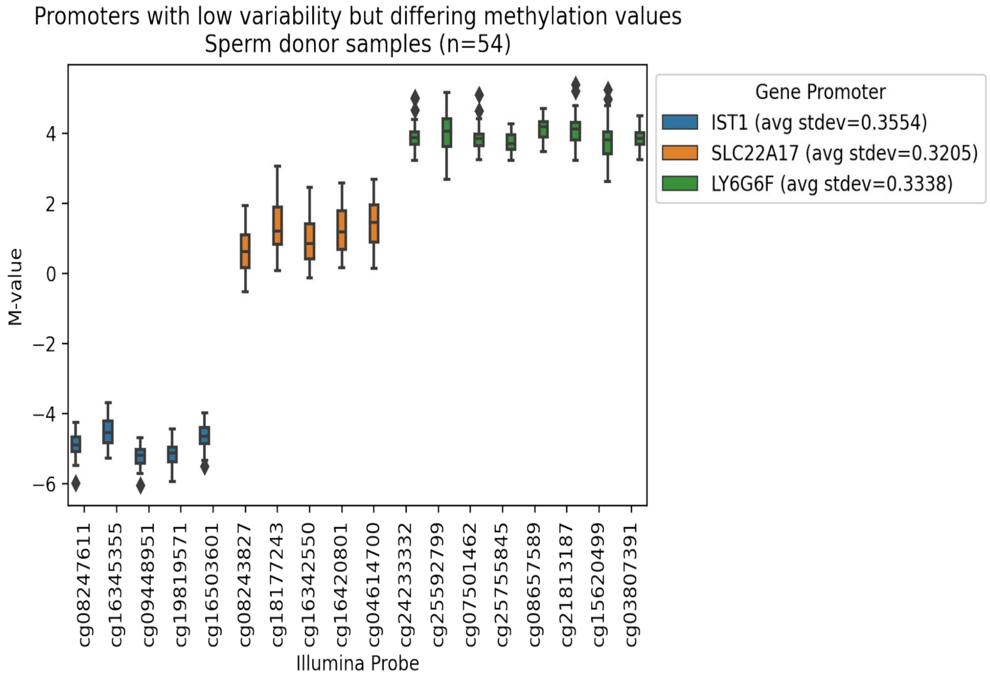


B

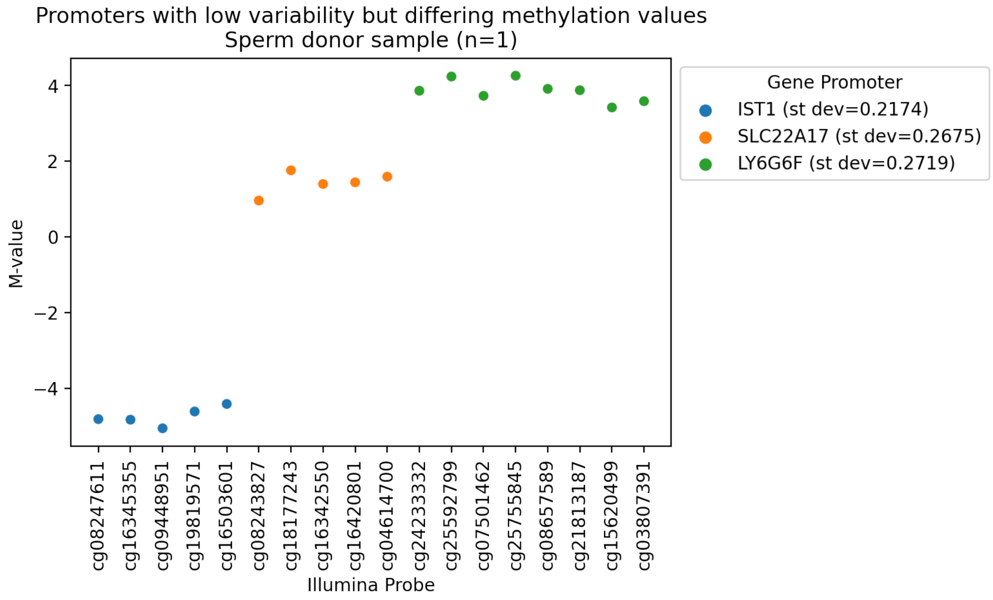

**Figure S6 | Principal component analysis of diseased and control samples**
This plot shows the principal component analysis of liver samples from healthy individuals and those with nonalcoholic fatty liver disease (NAFLD) and nonalcoholic steatohepatitis (NASH).

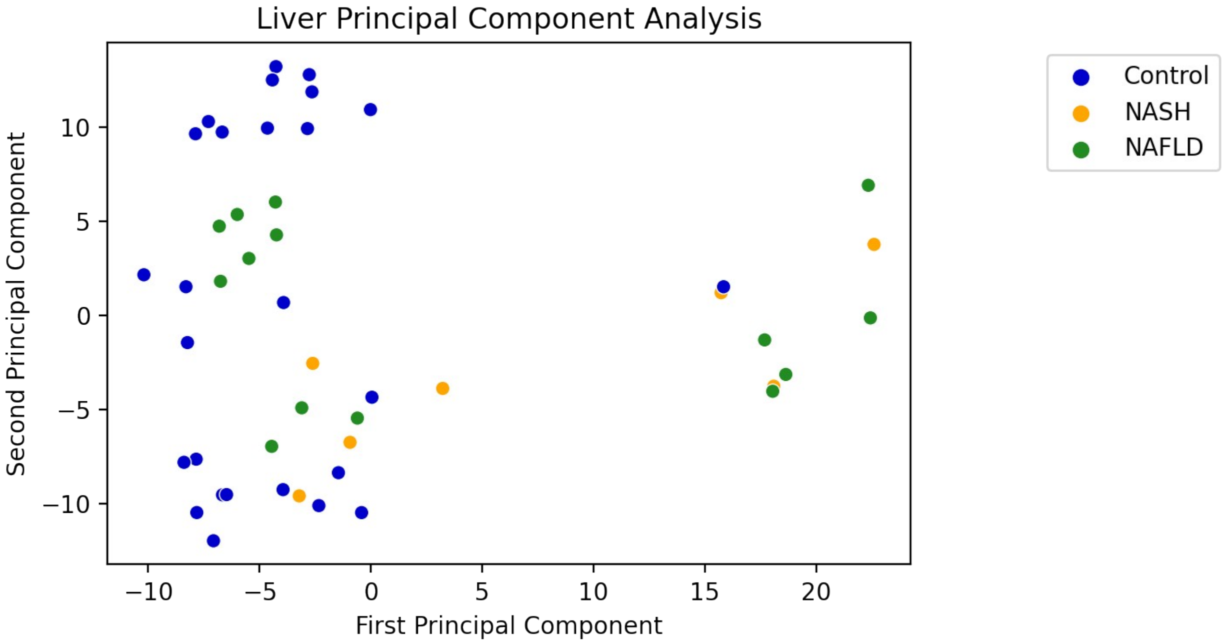

**Figure S7 | Hierarchical clustering of diseased and control tissue samples**
This plot shows the hierarchical clustering of diseased and control tissue samples. Clustering is based on the variability values of all promoters. (A) Hierarchical clustering of colon primary tumor samples and normal colon tissue samples (B) Hierarchical clustering of paired psoriatic skin lesion samples and normal skin samples (C) Hierarchical clustering of control, nonalcoholic fatty liver disease (NAFLD), and nonalcoholic steatohepatitis (NASH) liver samples

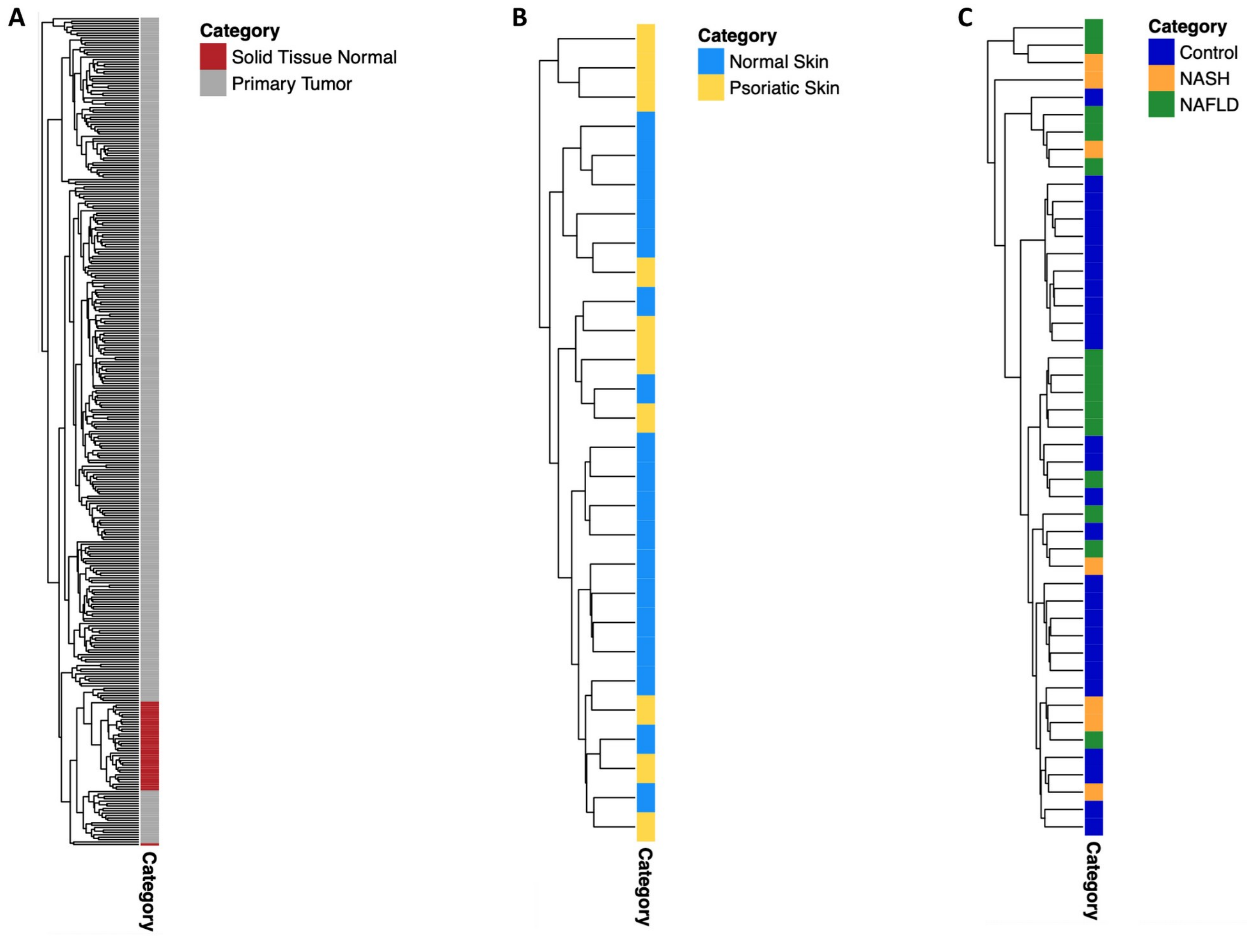

**Figure S8 | N-of-1 dysregulated promoter analysis of sperm samples from multiple cohorts**
These plots look at the number of dysregulated promoters from the most stable promoters in sperm. The “Sperm donor cohort (training)” is the cohort used to find the most stable promoters in sperm and set the promoter variability thresholds. Panel A shows the number of dysregulated promoters in various sperm sample cohorts on a linear scale and Panel B shows the same plot but on a logarithmic scale.

A

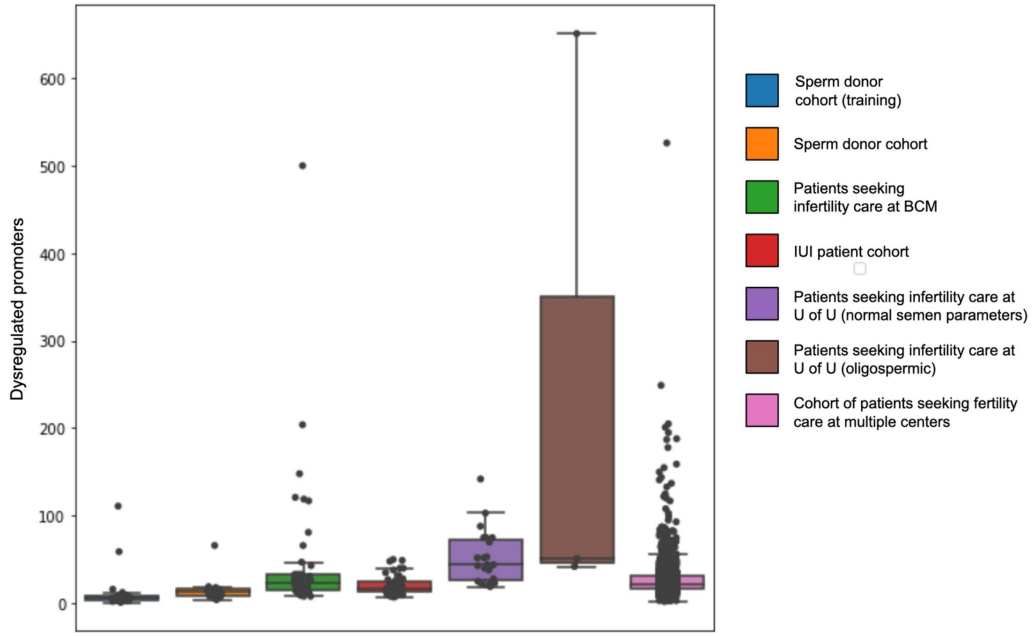


B

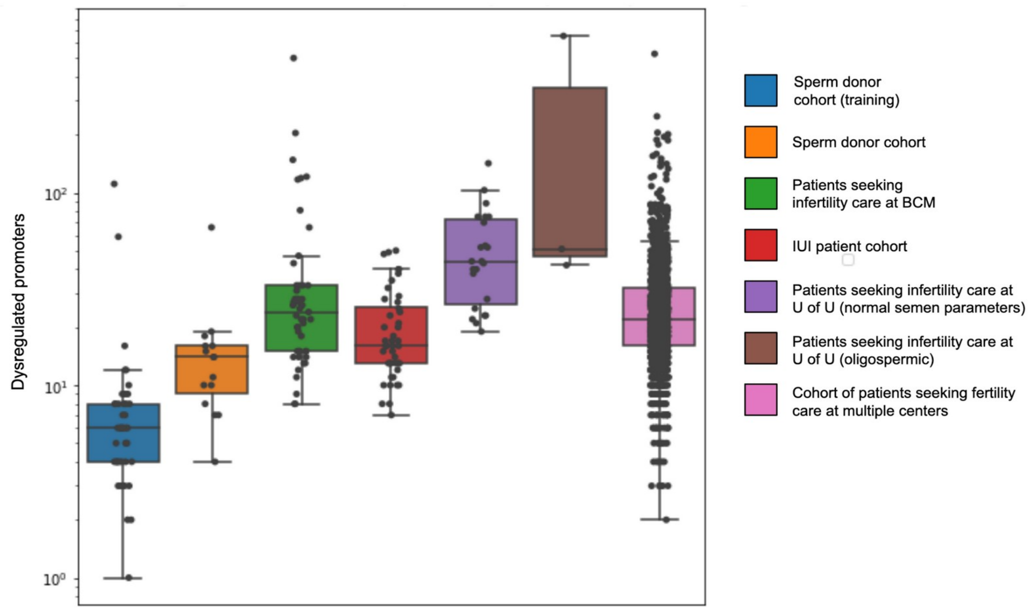

**Table S1 | Pregnancy/Birth rates in men with normal sperm concentration (≥ 15 million sperm / mL)**

| Variance Level | Pregnancy Rate from IUI | | Live Birth Rate from IUI | |
|---|---|---|---|---|
| Patients with the least number of dysregulated promoters (bottom 10th percentile) | 47.1% | p=0.082 | 39.2% | p=0.048* |
| Patients with the most number of dysregulated promoters (top 10th percentile) | 29.5% | | 20.5% | |

**Table S2 | Statistics from men with least and most number of dysregulated promoters**

| Variance Level | Pregnancy Rate from IUI | | Live Birth Rate from IUI | | Average / Median Age (Years) | | Average / Median Sperm Conc (M/mL) | | Average / Median BMI | |
|---|---|---|---|---|---|---|---|---|---|---|
| Patients with the least number of dysregulated promoters (bottom 10th percentile) | 48.1% | p=0.019* | 40.7% | p=0.011* | 33.3 / 33.5 | p=0.71 | 87.3 / 74.5 | p=0.89 | 29.7 / 28.7 | p=0.51 |
| Patients with the most number of dysregulated promoters (top 10th percentile) | 26.0% | | 18.0% | | 33.5 / 32.0 | | 86.1 / 58.3 | | 29.2 / 27.9 | |

**Table S3 | Ranking by sperm concentration**

| Concentration Level | Pregnancy Rate from IUI | | Live Birth Rate from IUI | |
|---|---|---|---|---|
| Patients with the highest sperm concentration (top 10th percentile) | 25.0% | p=0.52 | 23.1% | p=1.0 |
| Patients with the lowest sperm concentration (bottom 10th percentile) | 30.8% | | 23.1% | |
